# A novel transplantable model of lung cancer associated tissue loss and disrupted muscle regeneration

**DOI:** 10.1101/852442

**Authors:** Paige C Arneson, Alexandra M Ducharme, Jason D Doles

**Author notes:** These authors contributed equally to this work (Paige C Arneson, Alexandra M Ducharme).

## Abstract

**Background:** Cancer-associated muscle wasting (CAW), a symptom of cancer cachexia, is associated with approximately 20% of lung cancer deaths, and remains poorly characterized on a mechanistic level. Current animal models for lung cancer-associated cachexia are limited in that they: 1) primarily employ flank transplantation methods, 2) have short survival times not reflective of the patient condition, and 3) are typically performed in young mice not representative of mean patient age. This study investigates a new model for lung cancer-associated cachexia that can address these issues and also implicates muscle regeneration as a contributor to CAW.

**Methods:** We used tail vein injection as a method to introduce tumor cells that seed primarily in the lungs of mice. Body composition of tumor bearing mice was longitudinally tracked using NMR-based, echo magnetic resonance imaging (echoMRI). These data were combined with histological and molecular assessments of skeletal muscle to provide a complete analysis of muscle wasting.

**Results:** In this new lung CAW model we observed 1) progressive loss in whole body weight, 2) progressive loss of lean and fat mass, 3) a circulating cytokine/inflammatory profile similar to that seen in other models of CAW, 4) histological changes associated with muscle wasting, and 5) molecular changes in muscle that implicate suppression of muscle repair/regeneration. Finally, we show that survival can be extended without lessening CAW by titrating injected cell number.

**Conclusions:** Overall, this study describes a new model of CAW that could be useful for further studies of lung cancer-associated wasting and accompanying changes in the regenerative capacity of muscle. Additionally, this model addresses many recent concerns with existing models such as immunocompetence, tumor location, and survival time.

## BACKGROUND

Cancer cachexia is a complex syndrome associated with approximately 20% of lung cancer deaths; a hallmark symptom of cancer cachexia is cancer-associated muscle wasting (CAW) [1, 2]. Although this syndrome is associated with negative patient outcomes, it remains poorly understood on a mechanistic level. Many groups have investigated the inflammatory environment associated with CAW, in particular, inflammatory cytokines such as tumor necrosis factor alpha (TNFα) and interleukin 6 (IL6). Despite considerable time invested in this area, clinical trials targeting the inflammatory microenvironment alone have been unsuccessful [3, 4]. Other groups have focused on the proteolytic environment in muscle, as a potential driver of CAW, implicating both autophagy and ubiquitin-mediated pathways as potential drivers of CAW [5–7]. The current mechanistic understanding of CAW is largely based off of work done in mouse models, most notably the Lewis lung carcinoma (LLC) and C-26 models.

The LLC model of CAW relies on transplantation of LLC1 non-small cell lung cancer cells, derived from C57/B6 mice, into syngeneic, and thus immunocompetent, C57/B6 recipient mice. The LLC model is one of the few syngeneic and reproducible models for lung cancer/lung-cancer associated wasting available today [8]. Most commonly, LLC1 cells are implanted via flank injection, which results in large primary tumors around 7 days post implantation. A second popular model for CAW, the C26 model, was first described in 1990 and utilizes flank injection of colon 26 carcinoma cells [9]. Like the LLC model, this model displays elevated expression of proteasome components, which contribute to loss of muscle mass [10]. The C26 model presents with longer median survival (25 days) than the LLC model [10]. Notably, both the C26 and LLC models share a common weakness in that tumors do not arise in the tumor tissue of origin. Although recent studies have attempted to correct this issue using autochthonous genetically engineered mouse models (GEMMS), their utility and flexibility are limited. For example, in pancreatic cancer, there have been multiple efforts to create CAW models that focus on tumors arising in their native tissue via either genetic manipulation or orthotopic implantation [11–15]. The benefits and limitations of these models are addressed in more depth in the discussion. Despite these advances in the context of pancreas cancer, lung cancer models have lagged behind. Orthotopic transplantation has been established in rodent models, but requires an invasive surgery [16], [17]. Furthermore, many genetic methods to derive orthotopic tumors in lungs require viral delivery of Cre recombinase [18]. Complications arising in both these models are high and prompted us to develop a simple, reproducible, and scalable model of lung CAW.

We identified the need for better models of CAW that met the following criteria: 1) immunocompetent environment, 2) tumors arising in the appropriate/matching tissue of origin, and 3) ability to modulate the timing/duration of wasting. Here, we present studies on a novel mouse model for lung CAW, which has met these criteria. This model utilizes syngeneic, immunocompetent mice and tail-vein injection of cells as a modality for lung tumor seeding; shows a median survival of approximately 26 days that can be modulated as a function of injected cell number; and exhibits hallmarks of lung CAW. The model presented herein fills a gap in models for studying CAW, and will be a valuable resource for the wasting community moving forward.

## METHODS

### Animals

Mice were bred and housed according to NIH guidelines for the ethical treatment of animals in a pathogen-free facility at the Mayo Clinic (Rochester, MN campus). Mayo Clinic’s Institutional Animal Care and Use Committee (IACUC) approved all animal protocols. For 393P-induced CAW model, 150 μL 393P cells (containing 3 × 10^6^, 1 × 10^6^, or 3 × 10^5^ cells depending on group), or an equal volume of PBS (Gibco, 14190-144) (control) was injected via tail vein into 7-week-old male 129S2/SvPasCrl mice (Jackson Laboratories, Bar Harbor, ME). Development of cachexia was monitored twice weekly by body weight, fat, and lean mass. Mice were euthanized when endpoint criteria (weight loss greater than or equal to 20% of body weight, inability to ambulate, inability to reach food and/or water, tumors greater than or equal to 10% of body weight, tumors that have ulcerated, a body condition score of 1 or less using the IACUC approved scoring system) was reached, therefore, animals were euthanized on a case-by-case basis. For some animals their decline in health was rapid and resulted in spontaneous death. Survival time was reported as the time to reach endpoint criteria, or time until spontaneous death. Note that the criteria “tumors greater than or equal to 10% of body weight, tumors that have ulcerated” do not apply to primary lung tumors, but were used as assessment for palpable metastases that formed on the back and hindlimbs of some animals. Blood serum, heart muscle, lungs, tibialis anterior (TA), gastrocnemius (GR) and epididymal fat pad were collected immediately for analyses. Exact n for each experiment included in the figure legends.

### Animal imaging

Body weight, lean mass, and fat mass were measured twice weekly until endpoint. A baseline measurement was taken prior to start of study. Body composition was measured using an NMR-based echo magnetic resonance imaging EchoMRI-100/130 system (EchoMRI, Houston, TX, USA).

### Grip Strength Testing

Bioseb InVivo Research Instruments grip strength meter, model GT3, was used to measure grip strength of mice at their survival endpoint. Vehicle mice were assessed at the same time. We utilized the bar attachment for the grip strength meter. Each mouse had 3 total front limb grips, with resting time between each run. The 3 grips were averaged into a single data point for each mouse. Grips were not included in final count of 3 grips if the mouse used their hind limbs, gripped with only one paw, or refused to grip.

### Cell culture

Kras^LA1/+^;p53^R172H/Δg/+^ lung adenocarcinoma cell line (393P) was generated as previously described [19]. 393P cells were grown on tissue culture treated dishes in growth media consisting of DMEM media (Gibco #11995-065, lot 2051518) supplemented with 10% fetal bovine serum (Gibco #10099131, lot 2017488) and antibiotics. Cells were validated by IDEXX analytics and confirmed to be a pure culture and murine in origin. Cells were removed from plate with TrypLE (Gibco, 12604-013), counted, and resuspended in PBS prior to injection.

### Immunostaining

Left TA whole muscle was prepared for O.C.T embedding and cryosectioned. Tissue sections (8-10um) were post-fixed in 4% paraformaldehyde for 5 minutes at room temperature prior to immunostaining. Once fixed, isolated tissue sections were permeabilized with 0.5% Triton-X100 in PBS, followed by blocking with 3% BSA in PBS. Primary antibody incubations occurred at RT for 90 minutes or overnight in 4 degrees, and secondary antibody incubations followed at RT for 45 minutes in 3% BSA in PBS. The following antibodies were used in this study: Laminin (Sigma 4HB-2) and Pax7 (developmental hybridoma bank). Secondary antibodies were all Alexa fluorescent conjugates (488, 555, or 647) from Invitrogen or Jackson ImmunoResearch. Stained tissue sections were imaged on a Nikon Eclipse Ti-U camera and microscope system. Acquired images were analyzed for myofiber feret diameters using myovison and manual colocalization quantification using ImageJ.

### Cytokine Studies

Blood serum was collected from mice at endpoint. The MD31 cytokine panel was performed by Eve Technologies (Eve Technologies, Calgary, AB Canada) and samples were prepared as recommended by Eve Technologies. All samples were run in technical duplicate.

### Quantitative RT-PCR (qPCR)

Total gastrocnemius RNA was isolated, purified, and DNase I treated using Trizol and purified on RNeasy Mini kit columns (Qiagen, Mississauga, ON, Canada). RNA was quantified using a NanoDrop Spectrophotometer (ThermoScientific, Wilmington, DE, USA). Two μg of total RNA was reverse transcribed with primers using the High-Capacity cDNA Reverse Transcription Kit (Life Technologies, Carlsbad, CA, USA). QPCR was performed on a ViiA7 Quantitative PCR System (Applied Biosystems by Life Technologies, Austin, TX, USA). All samples were run in technical triplicate. Standard delta delta CT analysis was used post PCR. Primer sequences are available upon request.

### Lung and Heart Histopathology

At endpoint, lung and heart tissues were fixed in 4% paraformaldehyde. After 24 hours, tissue was moved to 70% ethanol. Lung tissue was embedded in paraffin and tissue sections (6 um) were incubated at 37 degrees for 60 minutes prior to staining with hematoxylin and eosin (H&E). Stained lung tissue sections were analyzed by GEMpath Inc. (Dr. Brad Bolon, Longmont, CO) and imaged on a Nikon Eclipse Ts2 microscope. Samples could be segregated with 100% reliability at using both the naked eye and the microscope. Heart tissue was embedded in paraffin and tissue sections (6 um) were incubated at 37 degrees for 60 minutes prior to post fixation in Bouin’s Solution, followed by staining with Picro Sirius Red. Stained tissue sections were imaged on a Motic EasyScan slide scanner and analyzed using ImageJ thresholding in the green channel.

### RNA Sequencing

RNA sequencing was done in collaboration with the Mayo Clinic Medical Genome Facility Genome Analysis Core and Mayo Clinic Bioinformatics Core. Transcripts with RPKM intensities equal to zero in all samples were removed and the remaining transcripts’ RPKM values were log2 transformed and used for subsequent analyses. Hierarchical clustering was performed using TIBCO Spotfire software. Clusters were formed by complete linkage clustering, and distances were measured by correlation. Principal component analysis (PCA)was performed using TIBCO Spotfire. RNAseq data set has been deposited in the Sequence Read Archive, as described in “Data Availability”.

### Statistics

Data are represented as the mean ± SD using GraphPad Prism (GraphPad Software, San Diego, CA) unless noted otherwise in the figure legends. Quantification of muscle cross sections using minimum feret diameter measurements was analyzed by non-linear regression (least squares method) and compared between conditions using an extra-sum-of-squares F test. All other comparisons between groups were performed using unpaired two-tailed student’s t tests or mixed-effects analysis, as noted in figure legends. For all analyses, a p<0.05 was considered significant (denoted with *)

### Data availability

The RNAseq dataset generated and analyzed during the current study is available in the Sequence Read Archive (SRA) (National Center for Biotechnology Information, NCBI), submission number SUB6913298, BioProject ID PRJNA604626. All other datasets generated during the current study are available from the corresponding author upon request.

## RESULTS

### Development of a transplantable model for lung cancer-associated tissue loss

Given the increasing need in the CAW field for models featuring tumors arising in their natural location, we sought to develop such a model in the context of lung cancer. To accomplish this, we used tail vein injection as a method whereby injected lung tumor cells implant primarily in the lungs [20]. We selected a cell line that was derived from Kras^LA1/+^;p53^R172H/Δg/+^ mice bearing lung adenocarcinomas[19]. Cells were cultured more than 10 passages prior to injection. For the injection, cells were suspended in PBS and control animals were injected with PBS vehicle only. Throughout the course of the study we assessed body composition via echoMRI until humane endpoint (see experimental schematic in **Figure 1A**). We performed survival studies on 3 cohorts of mice (total n=10 vehicle and 15 tumor) and observed significantly decreased survival for tumor bearing mice (average survival tumor=26.2 days) (**Figure 1B**). Upon macroscopic inspection of the lungs, tumor bearing mice had many small nodules, fibrotic tissue, and increased tissue density (**Figure 1C**). Upon microscopic inspection of the lungs in tumor-bearing mice, alveolar fields were replaced by multiple, coalescing tumor nodules (>70% in 3 animals, 40-50% in 2 animals assessed). The tumor nodules consisted of loosely packed, vacuolated cells and in some cases were associated with small clusters of intra-tumoral lymphocytes, usually at the margins of blood vessels or nodules. Lastly, the pleural surface was roughened by projecting fronds / plaques of tumor cells. In comparison, vehicle animals displayed expanded alveolar spaces and a smooth pleural surface (**Figure 1D**). Furthermore, we observed a statistically significant increase in total lung weight in tumor-bearing mice (**Figure 1E**). Due to the proximity of the lungs and heart, and an established literature base regarding cardiac cachexia, we assessed cardiac fibrosis [21]. Cardiac tissue stained with Picro Sirius Red did not show any difference in fibrosis between tumor-bearing and vehicle animals (**Supplemental Figure 1A-C**).

**Figure 1:**
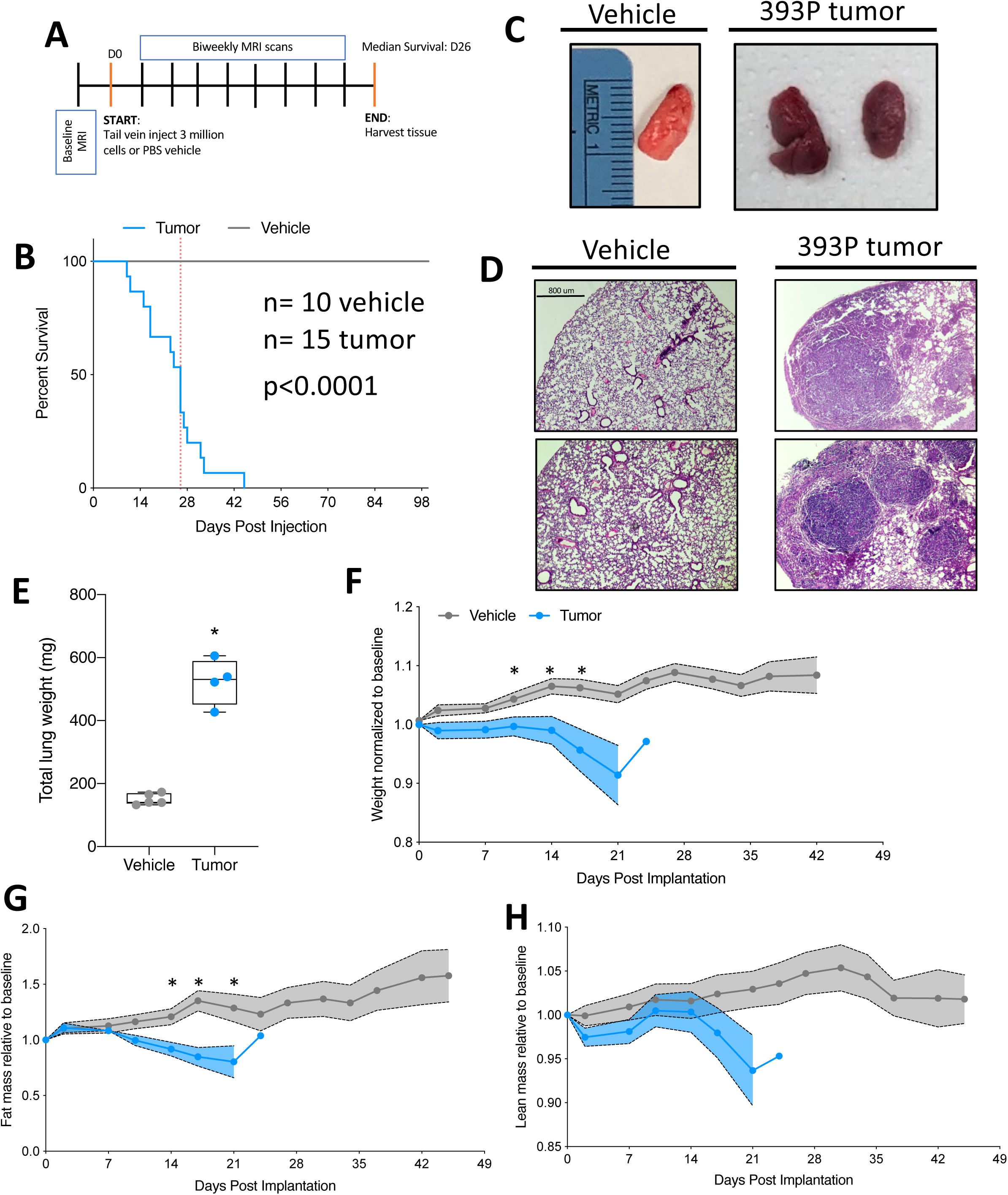
Transplantable model for lung cancer-associated tissue loss. **(A)** Schematic of the study design. D0=day tumor cells were injected into mice. All studies were survival studies, therefore, endpoint was variable. The median survival was 26 days. echoMRI scans were completed twice per week during the course of the study. **(B)** Survival curve for vehicle and tumor injected cells. Dotted vertical line indicates the median survival (26 days). n= 10 vehicle, 15 tumor. Data were compiled from 3 independent cohorts of animals. Survival is statistically different (p<0.0001) by Gehan-Breslow-Wilcoxon test. **(C**) Representative images of lungs from vehicle and tumor bearing mice. Ruler reference in centimeters. **(D)** Representative hematoxylin and eosin (H&E) staining of lung tissue from vehicle (left) and tumor bearing (right) mice. 2 representative images from 2 individual animals in each condition. Scale bar is 800 um. **(E)** Total lung weight (all lobes) from vehicle and tumor-bearing mice. Individual points represent individual animals. Boxes represent the inner quartiles and whiskers represent the minimum and maximum values. **(F)** Total mouse weight across study, normalized to pre-tumor baseline weight. Error bars are SEM. p values presented in the figure are the result of mixed-effects analysis, with Geisser-Greenhouse correction. Significance at individual points was determined by correction for multiple testing using false discovery rate (Benjamini and Hochberg). **(G)** echoMRI calculated total fat mass across study, normalized to pre-tumor baseline fat mass. Error bars are SEM, statistical analysis is as described above. **(H)** echoMRI calculated total lean mass across study, normalized to pre-tumor baseline lean mass. Error bars are SEM, statistical analysis is as described above. *p<0.05 by student’s t test n= 5 vehicle, 4 tumor-bearing 7-week-old male 129S2/SvPasCrl mice for all data except survival curve.

Concurrent with evaluations of tumor effects on overall survival, we performed longitudinal body composition analyses. First, we observed a significant difference in time and treatment co-variance for body weight, which was highlighted by significant decreases in body weight at 10, 14, and 17 days post tumor implantation (**Figure 1F**). Next, we observed a significant difference in time and treatment co-variance for total fat mass, which was highlighted by significant decreases in fat mass at 14, 17, and 21 days post tumor implantation (**Figure 1G**). Lastly, we observed a significant difference in time and group co-variance in total lean mass, although individual days had no significant difference (**Figure 1H**). Overall, there were decreases in total body weight, total fat mass, and total lean mass as the tumor progressed (the latter two variables measured using echoMRI) (**Figure 1F-H**).

### Lung cancer-associated molecular changes in serum and skeletal muscle

Loss in both body weight and lean mass are hallmarks of CAW. Many studies have implicated heightened inflammatory signaling as a driver of CAW [22–24]. We quantitatively assessed concentrations of 31 cytokines in the serum of tumor-bearing and vehicle animals (EveTechnologies) (n= 5 vehicle and 4 tumor). Ranked analysis (vertically by their average in the tumor-bearing animals; highest to lowest) revealed substantial differences between experimental groups (**Figure 2A**). Specifically, we observed significant differences in 4 factors: Eotaxin (down ~2 fold), C-X-C motif chemokine 5 (LIX, down ~10 fold), TNFα (up ~30 fold) and vascular endothelial growth factor (VEGF, up ~190 fold) (**Figure 2C**). Of note, TNFα accumulation has been previously implicated in CAW [22, 25].

**Figure 2:**
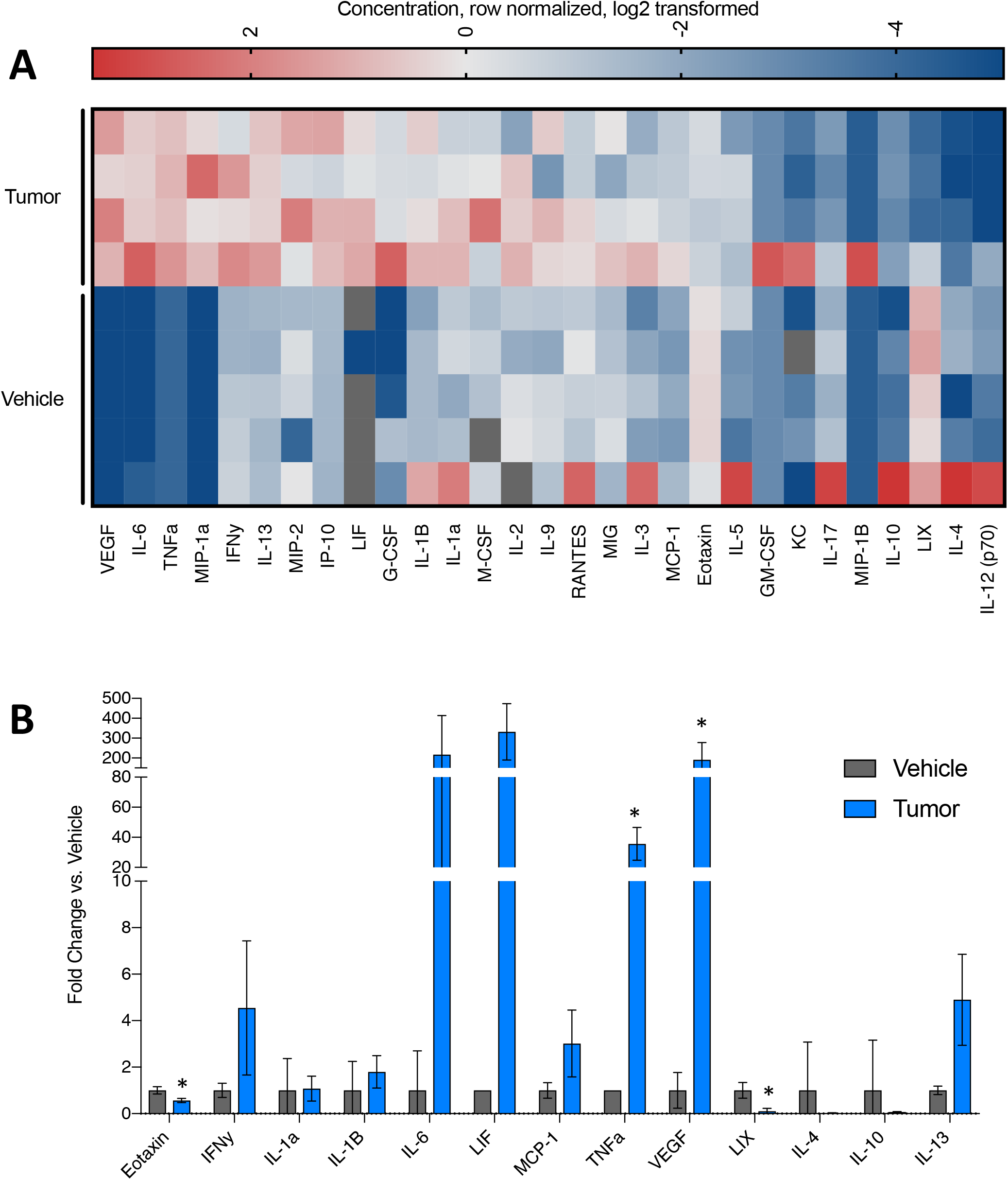
Lung cancer-associated inflammation. **(A)** Heat map of 29 cytokines profiled in the serum of vehicle or tumor-bearing mice. Features are sorted by the highest average of the tumor samples, from top to bottom. Values are the concentration, row normalized for each feature, then log2 transformed. Red= high expression, Blue= low expression. **(B)** Bar graph of cytokines commonly associated with muscle wasting and/or cancer cachexia. Individual points represent individual animals. Boxes represent the inner quartile and whiskers represent the minimum and maximum values. **(C)** Bar graph of cytokines significantly upregulated in tumor bearing animals. Individual points represent individual animals. Boxes represent the inner quartile and whiskers represent the minimum and maximum values. *p<0.05 in multiple t test with Holm-Sidak multiple testing correction. n= 5 vehicle, 4 tumor-bearing 7-week-old male 129S2/SvPasCrl mice.

In addition to serum cytokine profiling, we were interested in the transcriptional changes occurring in skeletal muscle, likely in response to these inflammatory changes. We therefore assessed the transcriptome of whole muscle from tumor-bearing and vehicle mice using RNA sequencing (RNAseq). Principle Component Analysis (PCA) and hierarchical clustering separated tumor-bearing from vehicle mice (**Figure 3A,B, Supplemental Figure 2**). We assessed differences in key pathways relevant to CAW such as muscle atrophy, autophagy, and hypoxia (**Figure 3C-E**). While we identified differential expression patterns in all three pathways, the most robust differences appeared to be in the muscle atrophy genes (**Figure 3C**).

**Figure 3:**
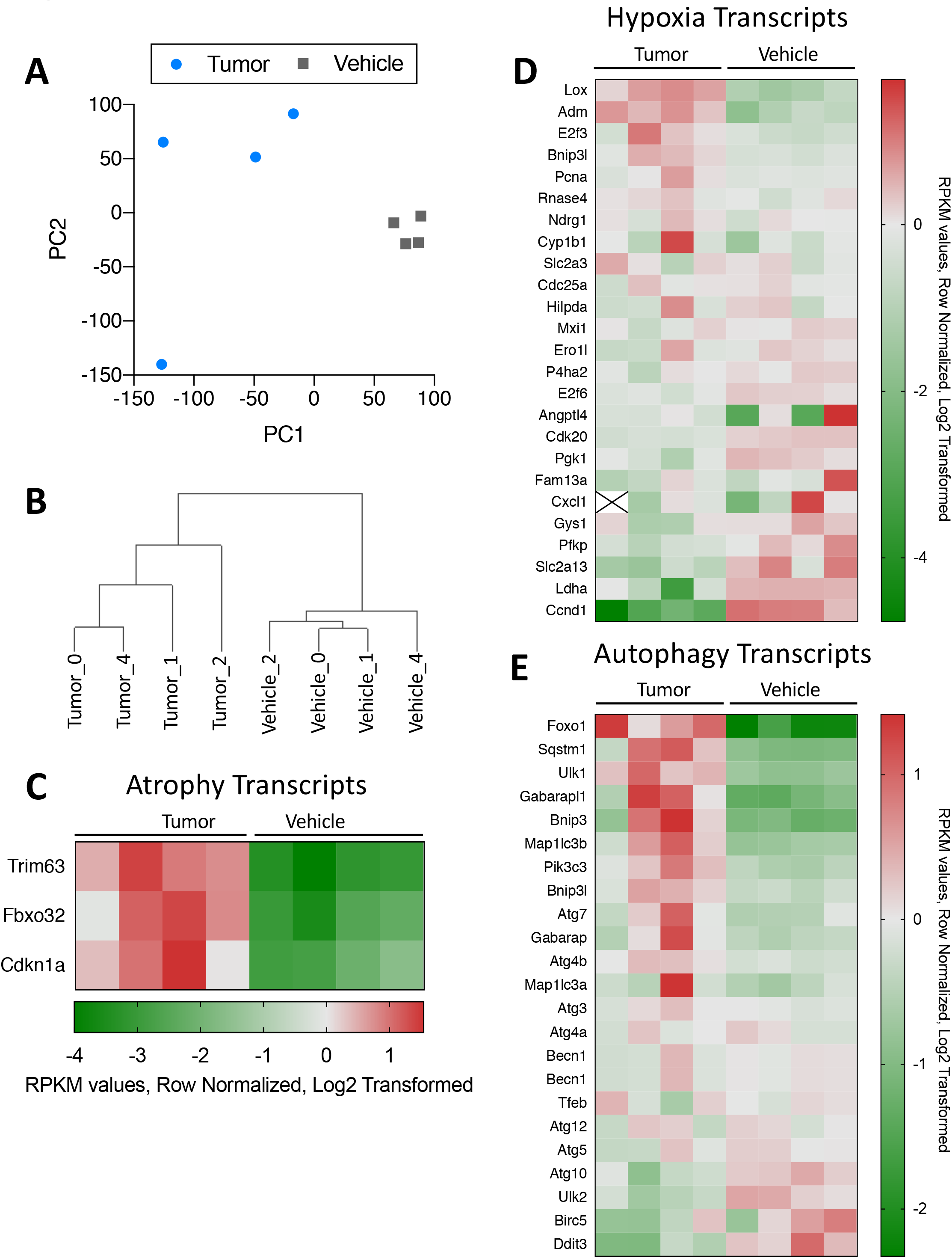
Lung cancer-associated transcriptional changes. **(A)** PCA analysis of all transcripts detected in tumor-bearing and vehicle mouse muscle. **(B)** Hierarchical clustering separates tumor-bearing and vehicle mouse muscle based on differential transcriptome patterns. **(C)** Heat map of selected transcripts commonly associated with muscle atrophy. Trim63 was significantly upregulated in tumor-bearing mouse muscle p=0.011. **(D)** Heat map of selected transcripts commonly associated with the hypoxia pathway. Of particular interest, 5 genes were significantly downregulated (Cdk20 p=0.0004, Ccnd1 p=0.0024, E2F6 p=0.0392) or upregulated (Adm p=0.0081, Lox p=0.0095). **(E)** Heat map of select4ed transcripts commonly associated with the autophagy pathway. There were not significant differences in expression of these genes. For all heat maps, features are sorted by the highest average of the tumor samples, from top to bottom. Values are the RPKM values, row normalized for each feature, then log2 transformed. Red = high expression, Green = low expression. Statistical significance was determined as p<0.05 in multiple t test with Holm-Sidak multiple testing correction. n= 4 vehicle, 4 tumor.

### Lung cancer-associated muscle loss

Considering both the loss of lean mass by echoMRI analysis, the accumulation of classical markers of CAW in the serum, and transcriptional changes associated with CAW, we next performed a histological and molecular assessment of skeletal muscle. First, we assessed tibialis anterior (TA) muscle cross sections by immunofluorescent staining for laminin, a marker of myofiber membranes, and DAPI (nuclei) (**Figure 4A**). Minimum feret diameters were measured, compiled, and calculated using MyoVision (mean feret diameter: 3 million = 40.4 um, vehicle = 44.4 um, and found a statistically significant decrease in the non-linear regressions fit to the histograms of minimum feret diameter distribution of tumor-bearing mice (**Figure 4B**) [26]. Additionally, we observed a significant decrease in TA weight in tumor bearing animals; this was not due to a reduction in the total number of myofibers in the TA of these animals (**Figure 4C-D**). This observed decrease in TA weight did not significantly affect grip strength at survival endpoints (**Supplemental Figure 3**) although tumor bearing mice were more likely to refuse to grip and were thus excluded from analysis (see Methods for analysis details/exclusion criteria). Since we observed a reduction in myofiber size, but not total number of myofibers, we postulated that this could be due to a decrease in the regenerative capacity of muscle. Using immunofluorescent staining, we queried the number of Pax7 positive muscle progenitor cells. We observed a statistically significant increase in Pax7 positive cells per millimeter squared of tissue (**Figure 4E**). Additionally, we assessed the number of centrally located nuclei (CLN), which mark actively regenerating myofibers containing recently fused muscle progenitor cells. We did not see any difference in the number of CLN per millimeter squared of tissue between vehicle and tumor-bearing mice (**Figure 4F**). Lastly, to acquire a molecular understanding of regenerative capacity of muscle from tumor-bearing animals, we used quantitative PCR to measure several transcripts associated with early myogenesis, late myogenesis/fusion, and muscle atrophy. Visualized in a heat map, we see a trend toward higher expression of atrophy transcripts (Trim63 and Fbxo32) and early makers of myogenesis (Pax7 and Myod1), and lower expression of late markers of myogenesis (**Figure 4G**).

**Figure 4:**
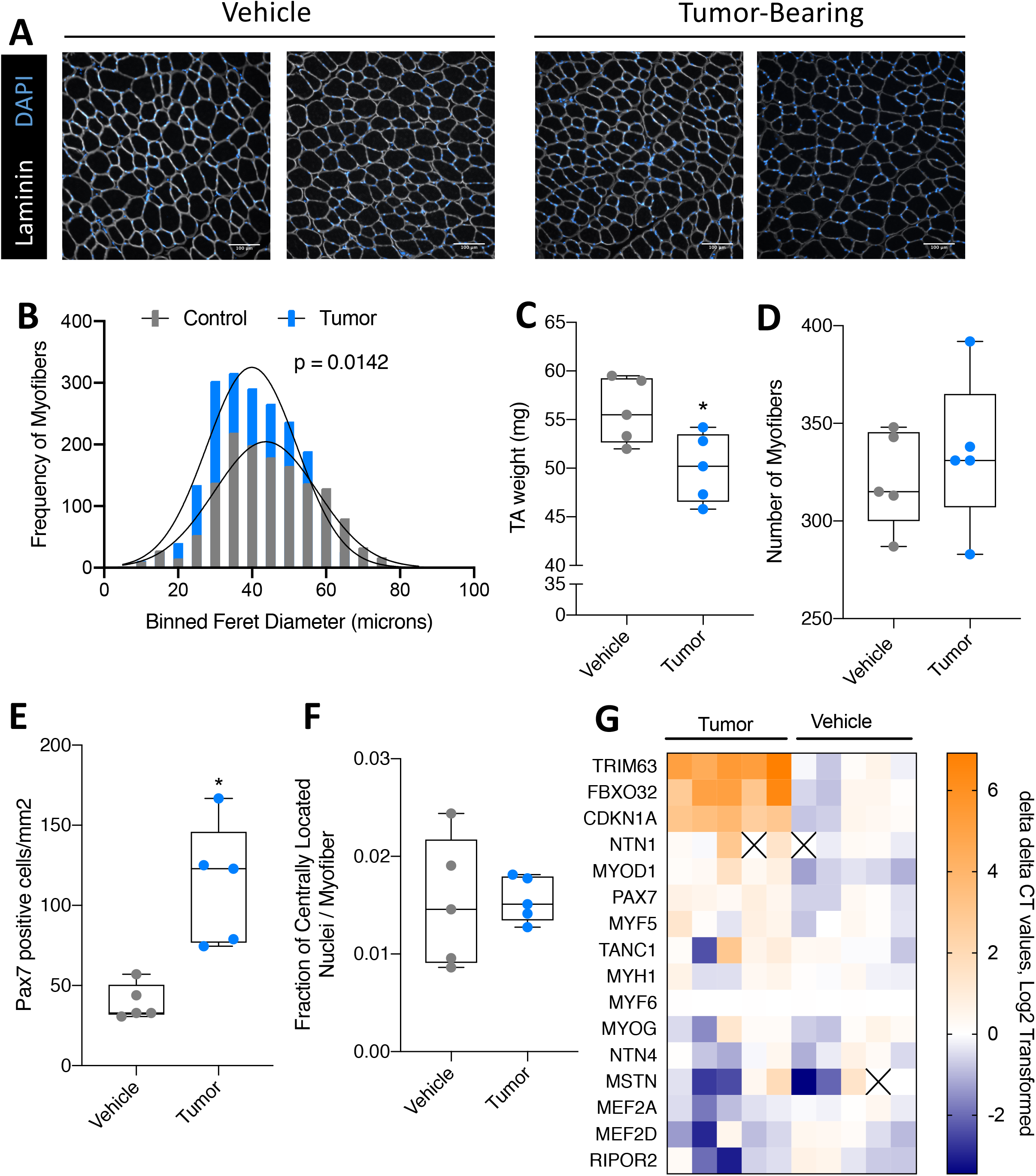
Lung cancer-associated muscle loss. **(A)** Representative images of vehicle (left two images) and tumor-bearing (right two images) tibialis anterior (TA) muscle cross sections. Immunofluorescent staining for Laminin (myofiber membranes, white) and DAPI (nuclei, blue). **(B)** Quantification of minimum feret diameters of myofibers from tumor-bearing and vehicle mice. Feret diameters were binned to a histogram and fit with a non-linear regression (gaussian, least squares regression). Myofibers of tumor-bearing animals were significantly smaller than vehicle; p= 0.0142. **(C)** Wet weight of TA muscle from vehicle and tumor-bearing animals. Individual points represent individual animals. Boxes represent the inner quartile and whiskers represent the minimum and maximum values. **(D)** Wet weight of gastrocnemius (GR) muscle from vehicle and tumorbearing animals. Individual points represent individual animals. Boxes represent the inner quartile and whiskers represent the minimum and maximum values. **(E)** Quantification of Pax7 positive cells per millimeter squared in TA cross sections of vehicle or tumor-bearing animals. Individual points represent individual animals. Boxes represent the inner quartile and whiskers represent the minimum and maximum values. **(F)** Quantification of centrally located nuclei per myofiber in TA cross sections. Individual points represent individual animals. Boxes represent the inner quartile and whiskers represent the minimum and maximum values. **(G)** Heat map depicting 16 transcripts assessed by qPCR in gastrocnemius muscle of vehicle or tumor-bearing animals. Transcripts are sorted by highest expression average of tumor sample from top to bottom. Orange= high expression, Blue= low expression. *P< 0.05 by student’s t test n= 5, 7-week-old male 129S2/SvPasCrl mice for each group.

### Transplanted cell number correlates with survival and CAW progression

Upon establishing that our lung cancer model induces loss in total body weight, lean mass, and is also associated with an impaired muscle regeneration phenotype, we sought to determine if survival time/CAW progression could be experimentally modulated. To accomplish this, we varied the number of cells injected as follows: 3 million cells (used in all previous figures), 1 million cells, 300,000 cells, and vehicle. We also tested 30,000 and 3,000 cells but found inconsistent inoculation of tumors and omitted these groups from further study (data not shown). We assessed survival of animals in each group and found that lowering the number of cells injected prolonged survival as evidenced by a statistical increase in survival from 3 million to 1 million cells (p=0.0208), 1 million to 300,000 cells (p=0.0024), and 300,000 cells to vehicle (p=0.0046) (**Figure 5A-B**). We also observed changes in lung condition, as evidenced by increased lung weight in 3 and 1 million injected cells, and macroscopic changes in lung condition similar to those observed previously (**Figure 5C-D**).

**Figure 5:**
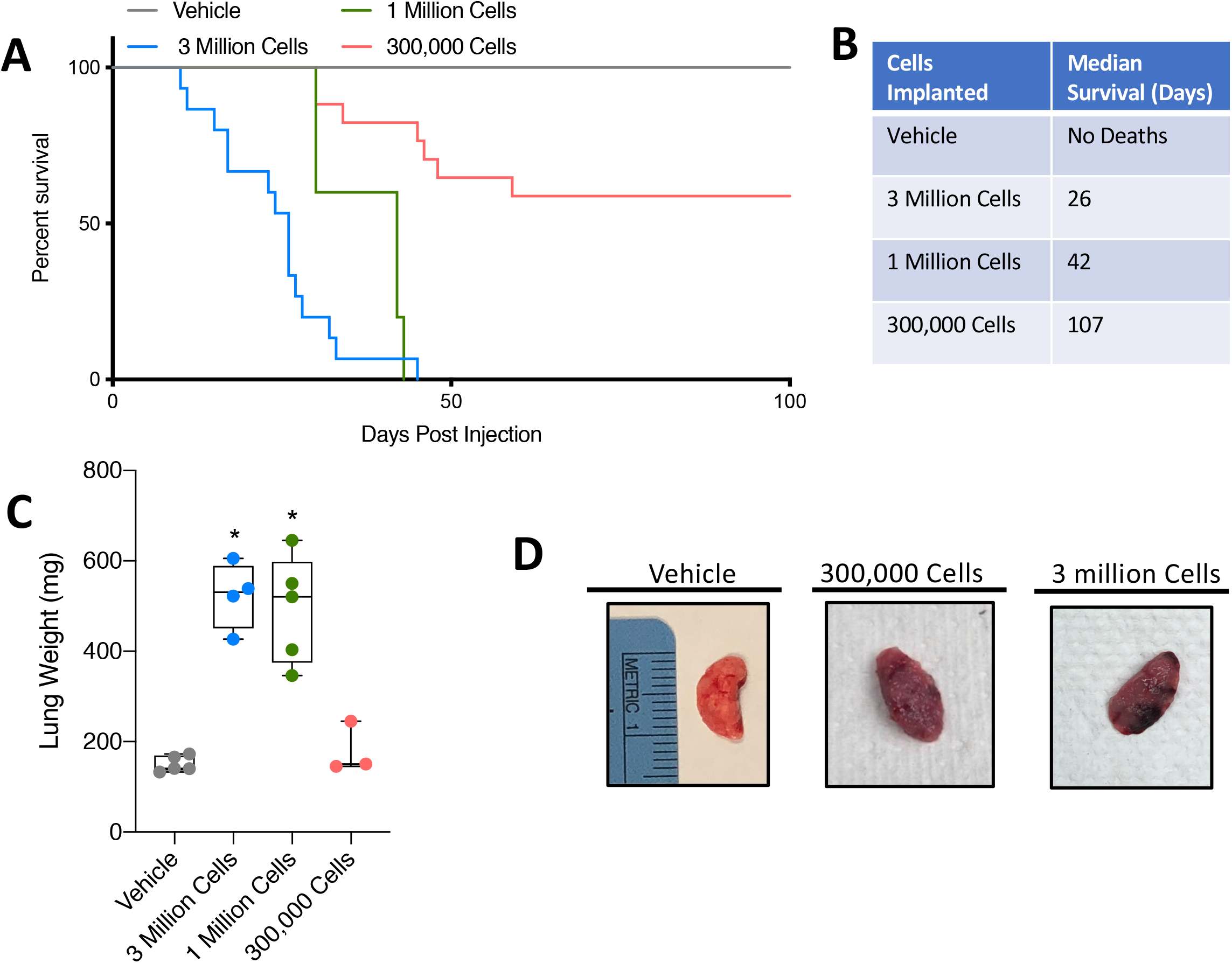
Tumor cell number titration modulates survival time. **(A)** Survival curve for mice injected with vehicle or a range of tumor cells (300,000, 1 million, or 3 million). **(B)** Table depicting the median survival for each group. **(C)** Total lung weight (all lobes) in milligrams for each group. Individual points represent individual animals. Boxes represent the inner quartile and whiskers represent the minimum and maximum values. *p<0.05 student’s t test. n= 5, vehicle; 4, 3-million; 5, 1-million; 3, 300,000 7-week-old male 129S2/SvPasCrl mice. **(D)** Representative images of lungs from vehicle, 300,000 cells and 3 million cells groups. Ruler reference is in centimeters.

We next performed body composition and muscle morphometrics analyses in mice receiving variable numbers of tumor cells. As before, we observed decreasing trends in body mass, lean mass and fat mass in tumor bearing mice of all groups (**Figure 6A-C**). Longitudinal weight, lean, and fat mass data for individual mice is available in **Supplemental Figure 4**. Specifically, we observed significant difference in time and treatment co-variance for body weight, in 3 million, 1 million, and 300,000 implanted cell groups compared to vehicle (**Figure 6A**). We also observed significant difference in time and treatment co-variance for total lean mass in 3 million and 1 million implanted cell groups compared to vehicle (**Figure 6B**). In lower implantation doses, we did not observe significant differences in total fat mass. Because tumors were restricted primarily to the lungs (with the exception of some metastases) (**Table 1**), it was not possible to discount tumor mass from echoMRI measurements. Evaluation of TA muscles from tumor-bearing mice revealed statistically decreases in TA mass in the 300,000 cells group, similar to the 3 million cells group (**Figure 6D**). Additionally, MyoVision-based assessment of minimum feret diameters revealed increasingly significant decreases as the cell injection number decreased (mean feret diameter: 1 million = 41.3 um, 300,000 = 32.6 um, vehicle = 44.4 um) (**Figure 6E**).

**Figure 6:**
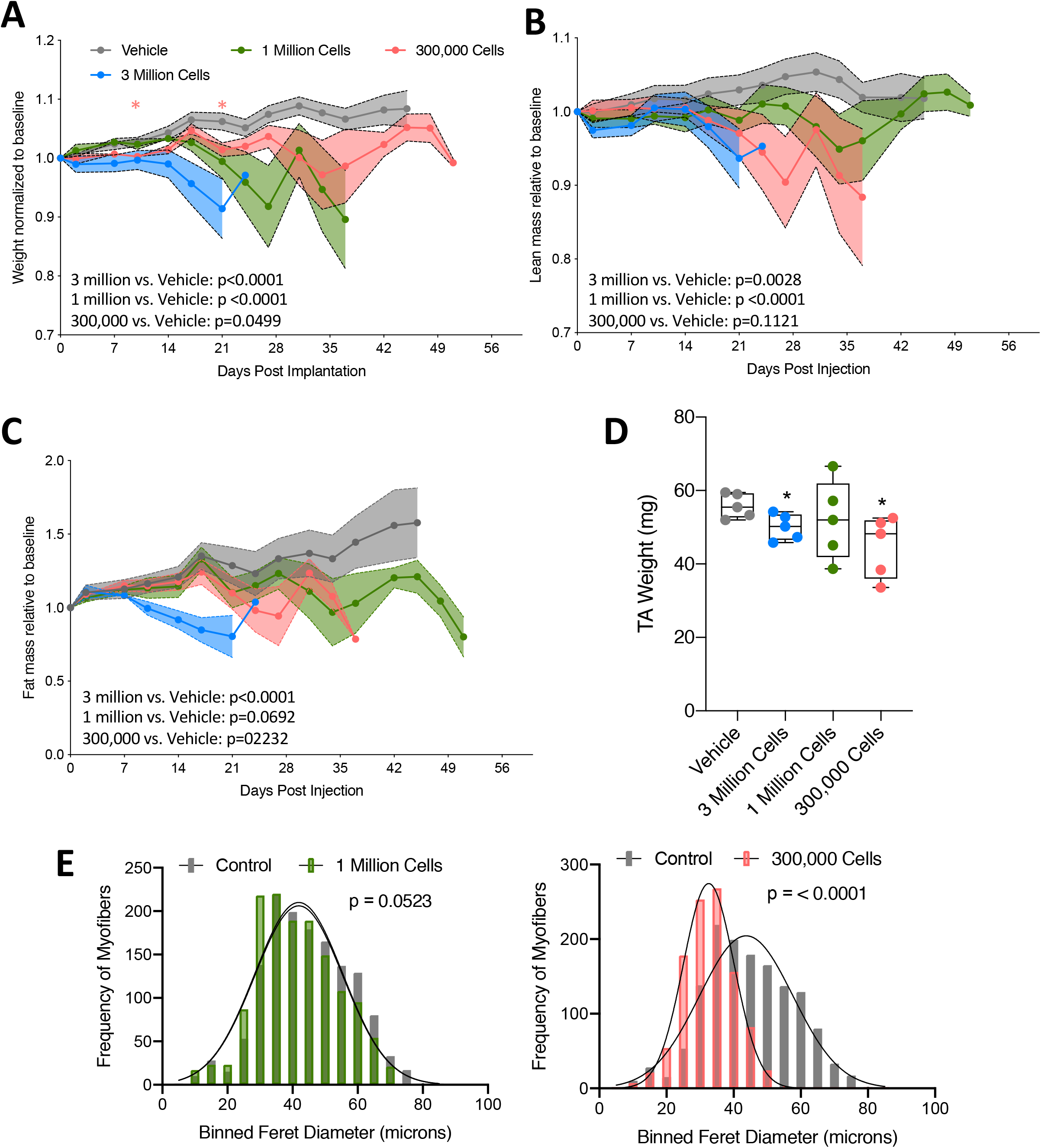
Tumor-bearing mice exhibit progressive tissue loss regardless of initial transplanted cell number. **(A)** Total mouse weight across study, normalized to pre-tumor baseline weight. Error bars are SEM, statistical analysis is as described in 1F. **(B)** echoMRI calculated total fat mass across study, normalized to pre-tumor baseline fat mass. Error bars are SEM, statistical analysis is as described in 1F. **(C)** echoMRI calculated total lean mass across study, normalized to pre-tumor baseline lean mass. Error bars are SEM, statistical analysis is as described in 1F. **(D)** TA weight in milligrams for each group in study. Individual points represent individual animals. Boxes represent the inner quartile and whiskers represent the minimum and maximum values. **(E)** Quantification of minimum feret diameters from muscles of tumor-bearing (1 million cells or 300,000 cells) and vehicle-treated mice. Feret diameters were binned to a histogram and fit with a non-linear regression (gaussian, least squares regression). Myofibers from tumor-bearing mice were smaller, p values calculated by extra sun-of-squares F test. 1 million vs. vehicle (left, p= 0.0523), 300,000 vs. vehicle (right, p<0.0001). n= 5, vehicle; 4, 3-million; 5, 1-million; 3, 300,000 7-week-old male 129S2/SvPasCrl mice.

**Table 1:** Metastatic characteristics of 393P lung cancer-associated wasting model. Percentage of animals bearing nodular tumors visible with the naked eye during necropsy and qualitative assessment of necropsied animals showing metastases in the thorax, lower back, and upper hindlimb regions.

| Group | % with Lung Tumor | % with Other Tumors | % with Metastasis in Thoracic Cavity |
| --- | --- | --- | --- |
| 3,000,000 (n=15) | 93 | 33 lower back<br>10 hindlimb | 73 |
| 1,000,000 (n=5) | 100 | 100 | 100 |
| 300,000 (n=15) | 67 | 33 lower back | 53 |
| Vehicle (n=10) | 0 | 0 | 0 |

## DISCUSSION

In order to gain a more thorough understanding of CAW, there is a need for models that more accurately reflect the patient condition. This new and novel model for CAW addresses several needs in the field such as: 1)immunocompetence (i.e. a syngeneic model), 2) exhibits both heightened catabolism and suppressed regeneration 3) exhibits both lean and fat mass loss, 4) exhibits an inflammatory microenvironment, 5) is flexible in timing and severity, 6) arises in the tumor organ of origin. To the last point, recent studies have highlighted the importance of tumor location on both tumor development and CAW [27–29]. Our model is an easy and effective method to induce tumor formation in the lungs and concomitant CAW, using lung adenocarcinoma cells.

With regard to points two through four, developing therapeutics for CAW is ultimately dependent on mouse models that reflect the complex molecular nature of this syndrome. We showed that this model exhibits an inflammatory signature similar to published signatures of CAW patients [22, 23, 30–32]. Our model has the added attraction that inflammatory status is not affected by tumor implantation, as our model does not require a major surgery or viral infection, known modulators of inflammatory status [33]. Additionally, our model supports numerous studies linking both elevated catabolism and impaired skeletal muscle regeneration as contributors to CAW [5–7, 34]. Our observation that there is an accumulation of Pax7+ cells in the TA muscle, but no change in the number of centrally located nuclei may be indicative of increased progenitor cell expansion, with failure to incorporate into the damaged/cachectic muscle. Although we probed genes related to fusion and progenitor cell maturation by qPCR and did not observe differences in expression, a more detailed assessment of the progenitor cell population would be necessary to make definitive conclusions about suppressed muscle regeneration in this model.

A major weakness of many CAW mouse models is the age of wasting onset. Genetic models, like the KPC model, are predisposed to spontaneously develop tumors, which gives investigators little control over the age of onset [13]. Additionally, disrupted muscle development may be a contributor to the loss of skeletal muscle mass in spontaneous genetic models, highlighting the difference between true atrophy and absence of growth. This principle also implicates transplantation models of CAW [35]. Although muscle development is typically considered complete at 3 months this anabolic state is unrepresentative of the patient population [13, 36, 37]. With cachectic cancers affecting a primarily elderly population, it is critical that new models are able to account for this confounding factor. For pancreatic cancer, by the KPP mouse has addressed this problem by expressing pancreas-specific oncogenes under Cre recombinase control [13]. However, for lung cancer, genetic models of CAW are only beginning to be explored [15]. Although our model has not yet been adapted for use in aged mice, it is amenable to assessing CAW in any age, including elderly (>18 month old) mice. This may lead to a better understanding of CAW mechanisms and more efficient translation to the clinic.

Another aspect of the patient condition poorly modeled in rodents is the severity and duration of wasting symptoms. Recently, it was established that cachexia syndrome could be divided into three clinical stages: pre-cachexia, cachexia, and refractory cachexia [38]. Although refractory cachexia is marked by a short (only three months) survival, wasting is present for much longer – an important observation that is difficult to study in rapidly progressing rodent CAW models. Importantly, with respect to intervention, some suggest that it is in the earlier stages (non-refractory) that therapeutics may be more effective [39]. Despite these features in patients, mouse models like LLC and C26 have rapid symptom onset and short survival times. Failing to capture the earlier stages of cachexia is likely a contributor to the failed clinical trials based on pre-clinical data derived using these models. The ability of our model to be titrated by cell dosage provides an important opportunity explore the timing and duration of wasting symptoms more thoroughly.

## CONCLUSIONS

Despite the many strengths highlighted in this study, most notable limitation is one common to all orthotopic lung cancer models, which is the difficulty for assessing tumor burden in living animals. This could be remediated by stably transfecting cells with a fluorescent reporter and utilizing *in vivo* imaging modalities. Additionally, in some cohorts, we observed high levels of metastasis outside the thoracic cavity. This may be representative of patients that exhibit metastases, but also complicates data interpretation. Nevertheless, the model presented here is an important step forward as CAW research progresses and will be a valuable resource for future research aimed at understanding the etiology of lung cancer-associated muscle and fat wasting.

## LIST OF ABBREVIATIONS

CAW: Cancer associated muscle wasting
LLC: Lewis lung carcinoma
CLN: Centrally-located nuclei
TA: Tibialis Anterior
qPCR: quantitative polymerase chain reaction
echoMRI: echo magnetic resonance imaging
H&E: Hematoxylin and Eosin
DAPI: 4′,6-diamidino-2-phenylindole
RNAseq: Ribonucleic acid sequencing
PCA: Principle Component Analysis
GR: Gastrocnemius
SEM: Standard Error of the Mean
SD: Standard Deviation
TNFα: Tumor Necrosis Factor Alpha
IL6: Interleukin 6
GEMMs: Genetically Engineered Mouse Models
IACUC: Institutional Animal Care and Use Committee
LIX: C-X-C Motif Ligand 5
VEGF: Vascular Endothelial Growth Factor
Trim63: Muscle RING Finger 1 (MuRF1)
Fbxo32: F-box Only Protein 32
Pax7: Paired Box 7
MyoD1: Myogenic Differentiation 1
NIH: National Institutes of Health
NIAMS: National Institute of Arthritis and Musculoskeletal and Skin Diseases
NCI: National Cancer Institute
RSTP: Regenerative Sciences Training Program

## DECLARATIONS

### Ethics approval and consent to participate

Mice were bred and housed according to NIH guidelines for the ethical treatment of animals in a pathogen-free facility at the Mayo Clinic (Rochester, MN campus). Mayo Clinic’s Institutional Animal Care and Use Committee (IACUC) approved all animal protocols.

### Consent for publication

Not Applicable

### Availability of data and materials

The datasets used during the current study are available from the corresponding author on reasonable request.

### Authors’ contributions

Study design: JDD, PCA, AMD. Data collection: PCA and AMD. Data analysis/interpretation: JDD, PCA, AMD. Writing and editing of manuscript: JDD, PCA, AMD. All authors approved this manuscript.

## Acknowledgements

The authors wish to thank members of the Doles lab for helpful discussions and manuscript suggestions, Dr. Yanan Yang for graciously giving our group the 393P cell line used for these studies, and Martina Gluscevic for her help with cardiac tissue histopathology assessments. Additionally, the authors extend thanks to Brad Bolon, DVM, MS, PhD of GEMpath INC for assisting in the pathological analyses. The authors also would like to acknowledge the following Mayo Clinic Core facilities that assisted with data collection: Medical Genome Facility Genome Analysis Core, and the Bioinformatics Core.

## REFERENCES

1. Zhu, R., et al., Updates on the pathogenesis of advanced lung cance-r induced cachexia. Thoracic cancer, 2019. 10(1): p. 8–16.

2. Baracos, V.E., et al., Body composition in patients with non-small cell lung cancer: a contemporary view of cancer cachexia with the use of computed tomography image analysis. Am J Clin Nutr, 2010. 91(4): p. 1133S–1137S.

3. Porporato, P.E., Understanding cachexia as a cancer metabolism syndrome. Oncogenesis, 2016. 5: p. e200.

4. Jatoi, A., et al., A placebo-controlled, double-blind trial of infliximab for cancer-associated weight loss in elderly and/or poor performance non-small cell lung cancer patients (N01C9). Lung Cancer, 2010. 68(2): p. 234–9.

5. Pettersen, K., et al., Cancer cachexia associates with a systemic autophagy-inducing activity mimicked by cancer cell-derived IL-6 transsignaling. Sci Rep, 2017. 7(1): p. 2046.

6. Yuan, L., et al., Muscle-specific E3 ubiquitin ligases are involved in muscle atrophy of cancer cachexia: an in vitro and in vivo study. Oncol Rep, 2015. 33(5): p. 2261–8.

7. Baracos, V.E., et al., Activation of the ATP-ubiquitin-proteasome pathway in skeletal muscle of cachectic rats bearing a hepatoma. Am J Physiol, 1995. 268(5 Pt 1): p. E996–1006.

8. Kellar, A., C. Egan, and D. Morris, Preclinical Murine Models for Lung Cancer: Clinical Trial Applications. Biomed Res Int, 2015. 2015: p. 621324.

9. Tanaka, Y., et al., Experimental cancer cachexia induced by transplantable colon 26 adenocarcinoma in mice. Cancer Res, 1990. 50(8): p. 2290–5.

10. Aulino, P., et al., Molecular, cellular and physiological characterization of the cancer cachexia-inducing C26 colon carcinoma in mouse. BMC Cancer, 2010. 10: p. 363.

11. Lewis, B.C., D.S. Klimstra, and H.E. Varmus, The c-myc and PyMT oncogenes induce different tumor types in a somatic mouse model for pancreatic cancer. Genes & development, 2003. 17(24): p. 3127–3138.

12. Shukla, S.K., et al., Silibinin-mediated metabolic reprogramming attenuates pancreatic cancer-induced cachexia and tumor growth.Oncotarget, 2015. 6(38): p. 41146.

13. Talbert, E.E., et al., Modeling Human Cancer-induced Cachexia. Cell Rep, 2019. 28(6): p. 1612–1622 e4.

14. Frese, K.K. and D.A. Tuveson, Maximizing mouse cancer models. Nature Reviews Cancer, 2007. 7(9): p. 654.

15. Goncalves, M.D., et al., Fenofibrate prevents skeletal muscle loss in mice with lung cancer. Proceedings of the National Academy of Sciences, 2018. 115(4): p. E743–E752.

16. Fujiwara, T., et al., Therapeutic Effect of a Retroviral Wild-Type p53 Expression Vector in an Orthotopic Lung Cancer Model. JNCI: Journal of the National Cancer Institute, 1994. 86(19): p. 1458–1462.

17. Howard, R.B., et al., Characterization of a highly metastatic, orthotopic lung cancer model in the nude rat. Clinical & experimental metastasis, 1999. 17(2): p. 157–162.

18. DuPage, M., A.L. Dooley, and T. Jacks, Conditional mouse lung cancer models using adenoviral or lentiviral delivery of Cre recombinase. Nat Protoc, 2009. 4(7): p. 1064–72.

19. Gibbons, D.L., et al., Contextual extracellular cues promote tumor cell EMT and metastasis by regulating miR-200 family expression. Genes Dev, 2009. 23(18): p. 2140–51.

20. Doles, J., et al., Suppression of Rev3, the catalytic subunit of Pol{zeta}, sensitizes drug-resistant lung tumors to chemotherapy. Proc Natl Acad Sci U S A, 2010. 107(48): p. 20786–91.

21. Okuda, J., et al., Persistent overexpression of phosphoglycerate mutase, a glycolytic enzyme, modifies energy metabolism and reduces stress resistance of heart in mice. PloS one, 2013. 8(8).

22. Ma, J.F., et al., STAT3 promotes IFNγ/TNFα-induced muscle wasting in an NF-κB-dependent and IL-6-independent manner. EMBO Mol Med, 2017. 9(5): p. 622–637.

23. Zimmers, T.A., M.L. Fishel, and A. Bonetto, STAT3 in the systemic inflammation of cancer cachexia.Semin Cell Dev Biol, 2016. 54: p. 28–41.

24. Hogan, K.A., et al., Tumor-derived cytokines impair myogenesis and alter the skeletal muscle immune microenvironment. Cytokine, 2017.

25. Magee, P., S. Pearson, and J. Allen, The omega-3 fatty acid, eicosapentaenoic acid (EPA), prevents the damaging effects of tumour necrosis factor (TNF)-alpha during murine skeletal muscle cell differentiation. Lipids Health Dis, 2008. 7: p. 24.

26. Wen, Y., et al., MyoVision: software for automated high-content analysis of skeletal muscle immunohistochemistry. J Appl Physiol (1985), 2018. 124(1): p. 40–51.

27. Muir, A., L.V. Danai, and M.G. Vander Heiden, Microenvironmental regulation of cancer cell metabolism: implications for experimental design and translational studies. Dis Model Mech, 2018. 11(8).

28. Yanagihara, K., et al., Development and characterization of a cancer cachexia model employing a rare human duodenal neuroendocrine carcinoma-originating cell line. Oncotarget, 2019. 10(25): p. 2435.

29. Erstad, D.J., et al., Orthotopic and heterotopic murine models of pancreatic cancer and their different responses to FOLFIRINOX chemotherapy. Disease models & mechanisms, 2018. 11(7): p. dmm034793.

30. Horikawa, N., et al., Expression of vascular endothelial growth factor in ovarian cancer inhibits tumor immunity through the accumulation of myeloid-derived suppressor cells. Clinical Cancer Research, 2017. 23(2): p. 587–599.

31. Tachibana, K., et al., IL-17 and VEGF are increased and correlated to systemic inflammation, immune suppression, and malnutrition in patients with breast cancer. European Journal of Inflammation, 2017. 15(3): p. 219–228.

32. Gelin, J., et al., Role of endogenous tumor necrosis factor alpha and interleukin 1 for experimental tumor growth and the development of cancer cachexia. Cancer Res, 1991. 51(1): p. 415–21.

33. Pfeifer, A. and I.M. Verma, Gene therapy: promises and problems. Annual review of genomics and human genetics, 2001. 2(1): p. 177–211.

34. He, W.A., et al., NF-κB-mediated Pax7 dysregulation in the muscle microenvironment promotes cancer cachexia. J Clin Invest, 2013. 123(11): p. 4821–35.

35. White, R.B., et al., Dynamics of muscle fibre growth during postnatal mouse development. BMC developmental biology, 2010. 10(1): p. 21.

36. Aoki, T., et al., Evolution of peripheral lung adenocarcinomas: CT findings correlated with histology and tumor doubling time. American journal of roentgenology, 2000. 174(3): p. 763–768.

37. Zhang, Y., et al., Frequency of driver mutations in lung adenocarcinoma from female never-smokers varies with histologic subtypes and age at diagnosis. Clinical cancer research, 2012. 18(7): p. 1947–1953.

38. Fearon, K., J. Arends, and V. Baracos, Understanding the mechanisms and treatment options in cancer cachexia. Nat Rev Clin Oncol, 2013. 10(2): p. 90–9.

39. Fearon, K., et al., Definition and classification of cancer cachexia: an international consensus. Lancet Oncol, 2011. 12(5): p. 489–95.

