## Supplemental Data for "A novel transplantable model of lung cancer associated tissue loss and disrupted muscle regeneration"

*Corresponding author and lead contact

**SUPPLEMENTAL DATA**

Supplemental Figure 1

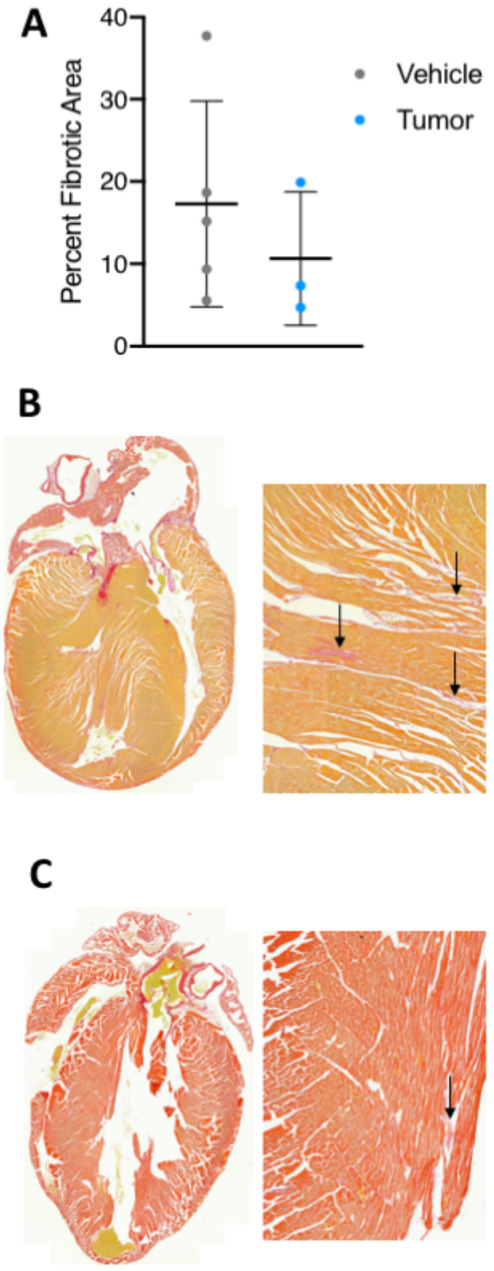


**Supplemental Figure 1: Assessment of Cardiac fibrosis. (A)** Comparison of percent fibrotic areas in whole heart longitudinal sections from vehicle and tumor-bearing mice. There was no significant difference in percent fibrotic area. **(B)** Representative images of a heart longitudinal section (left), and zoomed in area of fibrosis (right) from a tumor-bearing mouse. **(C)** Representative images of a heart longitudinal section (left), and zoomed in area of fibrosis (right) from a vehicle mouse. Black arrows are marking areas of positive staining for fibrosis.

### Supplemental Figure 2

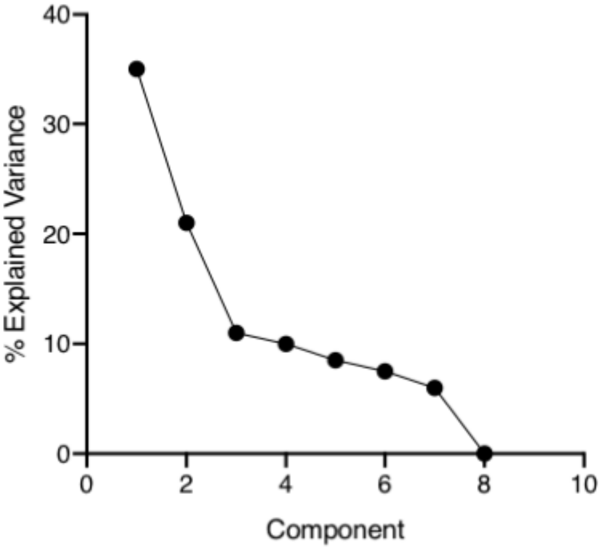


**Supplemental Figure 2: Explained variance for the principle component analysis (PCA).** PCA analysis and explained variance were generated in TIBCO Spotfire.

### Supplemental Figure 3

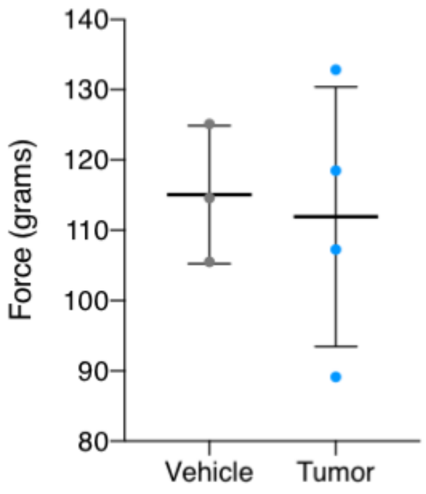


**Supplemental Figure 3: Grip strength at survival endpoint.** Grip strength was tested for tumor-bearing and vehicle mice at the survival endpoint for the tumor-bearing mice (vehicle mice were also sacrificed at this point). Each mouse had 3 grips that were averaged into one data point plotted above. Each data point plotted corresponds to one mouse. No statistical significance was found between groups. N=3 vehicle, 4 tumor-bearing.

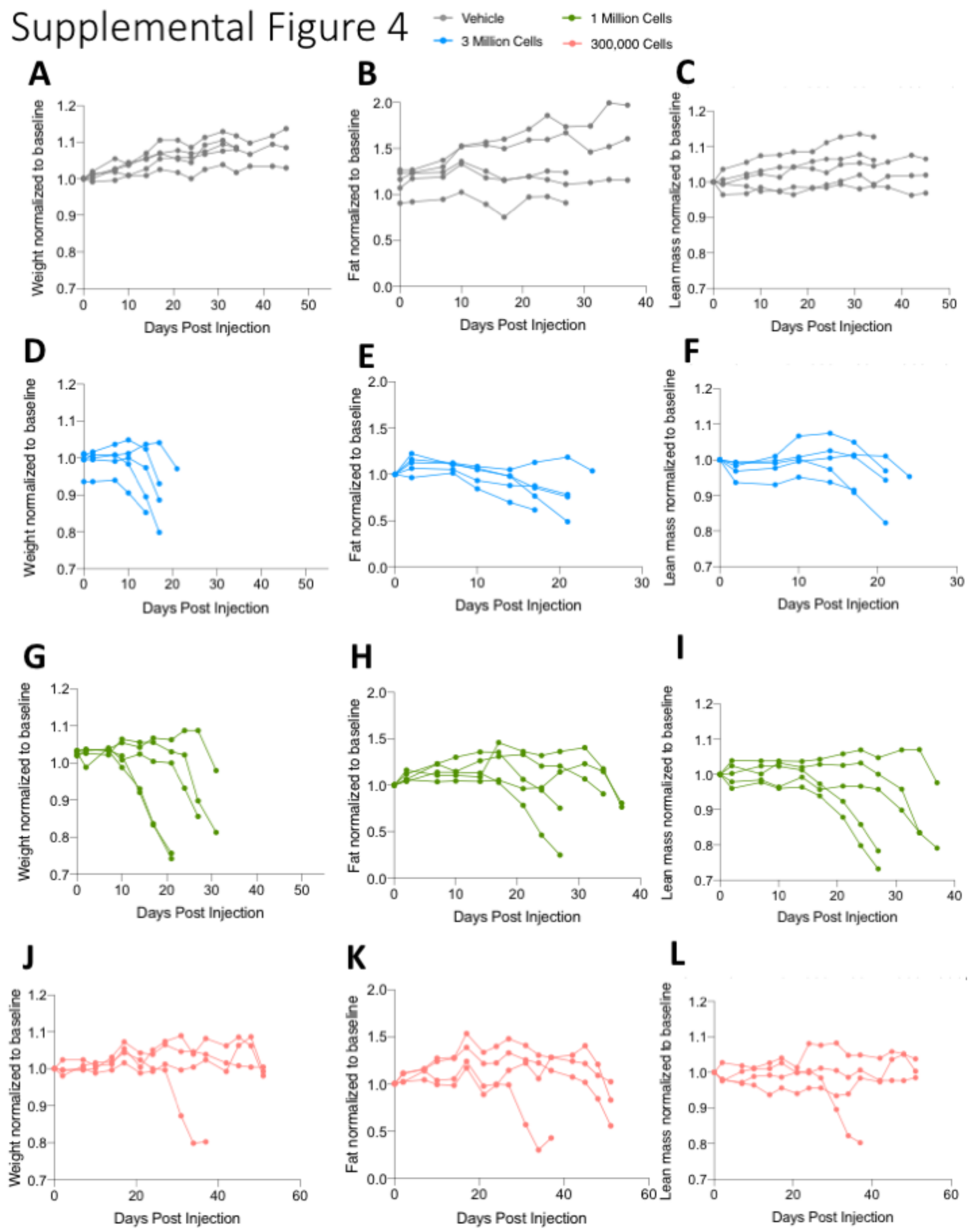


**Supplemental Figure 4: Longitudinal body composition assessment for individual mice.** Total mouse weight across study, normalized to pre-tumor baseline weight. Each line represents an individual animal in the following groups: Vehicle **(A)**, 3 million injected cells **(D)**, 1 million injected cells **(G)**, 300,000 injected cells **(J)**. echoMRI calculated total fat mass across study, normalized to pre-tumor baseline fat mass. Each line represents an individual animal in the following groups: Vehicle **(B)**, 3 million injected cells **(E)**, 1 million injected cells **(H)**, 300,000 injected cells **(K)**. echoMRI calculated total lean mass across study, normalized to pre-tumor baseline lean mass. Each line represents an individual animal in the following groups: Vehicle **(C)**, 3 million injected cells **(F)**, 1 million injected cells **(I)**, 300,000 injected cells **(L)**.  n= 5 vehicle, 3 million injected cells, and 1 million injected cells; 300,000 injected cells. 7-week-old male 129S2/SvPasCrl mice for all groups.
